# Pathway based factor analysis of gene expression data produces highly heritable phenotypes that associate with age

**DOI:** 10.1101/016154

**Authors:** Andrew Brown, Zhihao Ding, Ana Viñuela, Dan Glass, Leopold Parts, Tim Spector, John Winn, Richard Durbin

## Abstract

Statistical factor analysis methods have previously been used to remove noise components from high dimensional data prior to genetic association mapping, and in a guided fashion to summarise biologically relevant sources of variation. Here we show how the derived factors summarising pathway expression can be used to analyse the relationships between expression, heritability and ageing. We used skin gene expression data from 647 twins from the MuTHER Consortium and applied factor analysis to concisely summarise patterns of gene expression, both to remove broad confounding influences and to produce concise pathway-level phenotypes. We derived 930 “pathway phenotypes” which summarised patterns of variation across 186 KEGG pathways (five phenotypes per pathway). We identified 69 significant associations of age with phenotype from 57 distinct KEGG pathways at a stringent Bonferroni threshold (*P* < 5.38 × 10^−5^). These phenotypes are more heritable (*h*^2^ = 0.32) than gene expression levels. On average, expression levels of 16% of genes within these pathways are associated with age. Several significant pathways relate to metabolising sugars and fatty acids, others with insulin signalling. We have demonstrated that factor analysis methods combined with biological knowledge can produce more reliable phenotypes with less stochastic noise than the individual gene expression levels, which increases our power to discover biologically relevant associations. These phenotypes could also be applied to discover associations with other environmental factors.

## 1. Introduction

Ageing is a multifactorial process, reflecting how the physical state of an organism accumulates changes. Amongst these, we observe changes in gene expression. Microarrays and more recent RNA-seq technologies allow the simultaneous quantification of cell population average mRNA abundance for thousands of genes. In the case of ageing, consistent patterns of age-related changes in gene expression have been observed across several tissues and species [Lu et al., 2004], such as over-expression of inflammation and immune-response genes and under-expression of genes involved in energy metabolism in older samples [de Magalhaes et al., 2009]. Given this commonality of function amongst genes which show age related changes in expression, we decided to investigate ageing dependent gene expression in the context of biological knowledge on the function of genes, as provided by pathway annotations.

Array expression experiments generate high dimensional structured data sets, in which there are correlated patterns across large numbers of genes. Some of these are due to known technical or biological effects such as batch effects and cell growth stage, which when not the focus of the analysis can be removed by fitting them as covariates. However, even after this, there is typically substantial structural correlation. In previous studies, these can be represented by linear components of expression measurements, or factors, that can be inferred using methods such as principal components analysis (PCA) or factor analysis [Leek and Storey, 2007, Parts et al., 2011]. When the aim is to discover local effects, such as cis genetic regulation, the resulting factors can be treated as nuisance variables and removed from further analysis. This has been seen to increase power in analysis [Pickrell et al., 2010]. Conversely, if the aim is to differentiate between a case and control condition using expression, then factors viewed as global phenotypes could be more effective classifiers than local phenotypes [Hastie et al., 2000].

Recently we applied factor analysis methods in a two stage procedure to generate phenotypes representing expressions of groups of genes [Stegle et al., 2012]. After regressing out global factors, as in Parts et al. [2011], expression levels for groups of functionally related genes, as defined by annotations from pathway databases, were treated as new expression datasets and the same factor analysis methods were used to construct pathway factors. The factors constructed on pathway sets of genes were taken as concise summaries of common expression variation across each pathway. We test these factor values below as phenotypes, and so refer to them as phenotype factors or, in some cases, just phenotypes.

Here, we apply this method to gene expression data from abdominal skin tissues from 647 samples. Unlike previous studies which have concentrated on genetic variants which regulate multiple genes within a pathway [Stegle et al., 2012], we focus here on discovering associations between gene expression and age. We obtain our pathway gene sets from the Kyoto Encyclopedia of Genes and Genomes (KEGG) pathways [Kanehisa et al., 2004]. Subsequently, by looking for associations between these new pathway phenotypes and age, we discover groups of functionally related genes with a common response to ageing which can be used as biomarkers describing molecular changes with age.

With data from a twin cohort containing both monozygotic and dizygotic twins, we can estimate proportions of variance explained by age, genetic variation, common environmental variation, and unique environmental variation (noise). Stochasticity in gene expression, which will form part of the unique environment component, is believed to play a role in the ageing process [Bahar et al., 2006]. By investigating sources of variation within the pathway phenotypes, we find that they are more robust than the expression of individual genes, with less unique environment variation. This explains some of our success at discovering associations with age.

## 2. Methods

### 2.1 Expression profiling

The data analysed here are part of the MuTHER project (Multiple Tissue Human Expression Resource - http://www.muther.ac.uk/, [Nica et al., 2011]) and were downloaded from the ArrayExpress archive, accession no. E-TABM-1140. In summary, the study included 856 Caucasian female individuals (336 monozygotic (MZ) and 520 dizygotic (DZ) twins) recruited from the TwinsUK Adult twin registry [Moayyeri et al., 2012]. The age at sampling ranged from 39 to 85 years with a mean age of 59 years. Punch biopsies (8mm) were taken from relatively photo-protected infra-umbilical skin. Subcutaneous adipose tissue was dissected from each biopsy and the remaining skin tissue was weighed and stored in liquid nitrogen. Expression profiling of this skin tissue was performed using Illumina Human HT-12 V3 BeadChips where 200ng of total RNA was processed according to the protocol supplied by Illumina. All samples were randomised prior to array hybridisation and the technical replicates were always hybridised on different beadchips. Raw data were imported to the Illumina Beadstudio software and probes with fewer than three beads present were excluded. Log2-transformed expression signals were then normalised separately per tissue with quantile normalisation of the replicates of each individual followed by quantile normalisation across all individuals as previously described [Grundberg et al., 2012]. Post-QC expression profiles were subsequently obtained for 647 individuals. The Illumina probe annotations were cross-checked by mapping the probe sequence to the NCBI Build 36 genome with MAQ [Li et al., 2008]. Only uniquely mapping probes with no mismatches and either an Ensembl or RefSeq ID were kept for analysis. Probes mapping to genes of uncertain function (LOC symbols) and those encompassing a common SNP (1000G release June 2010) were further excluded, leaving 23,555 probes used in the analysis.

##### Box 1: Modelling

We model phenotype *y*_*i*_ of individual *i* (age *Ai*) as follows:

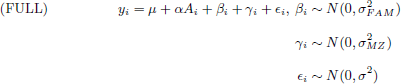

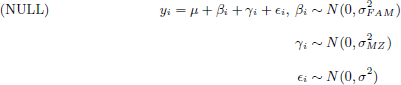

To correctly model the twin structure we enforce that β_i_ = β_j_ when i and j are twins, and γ_i_ = γ_j_ when i and j are monozygotic twins (capturing the increased genetic correlation of monozygotic twins).

From the full model we can define heritability (*h*^2^), proportion of environmental variance explained by age (*ρ*_*a*_) and the proportion of variance explained by the unique environment (*ρ*_*e*_) as:

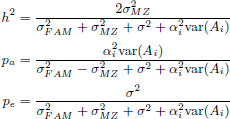

### 2.2 Gene expression pathway factors

In a two step approach, factor analysis methods were first used to discover patterns of common variation across the entire dataset. The software package PEER [Parts et al., 2011] was applied using the default settings and using technical measurements (experimental batch, RNA quality and concentration) as covariates to create 5 global factors, which in total explained 35.7% of the variation in the dataset. For each individual, a factor is a weighted sum of all the gene expression measurements of that individual. The weights are chosen so that the factors iteratively explain the maximum amount of variation in the dataset subject to certain prior assumptions; these factors produce concise summaries of consistent patterns of expression for large numbers of genes.

We then used KEGG pathway annotation (186 pathways) as prior information to group genes into pathways. This allows inference of PEER factors for each pathway that we refer to as phenotype factors, in contrast to the global factors previously described. As before, these factors are weighted sums of gene expression measurements, but in this case only of genes within the pathway. Since global factors have been removed from the dataset prior to calculation of phenotype factors, these factors are unlikely to capture global effects on gene expression, but instead pathway specific patterns of expression. If a large enough module of genes within the pathway is co-expressed then one factor will capture the same pattern of co-expression across individuals. Equally, groups of genes could show opposite patterns of expression; this antagonistic gene expression can also be reflected as a factor value which correlates across individuals with one set of genes and is anti-correlated with the other set of genes. Individual genes can contribute positively or negatively to the weighted sum (indicated by the sign of the corresponding weight), meaning that a positive correlation between age and phenotype factor can be induced by negative correlations with individual genes.

We grouped the expression data set into 186 pathway subsets. For each pathway we created five pathway phenotypes using PEER with the default settings. We consider the learnt pathway factor values across individuals as five new phenotypes which can be investigated for associations with age. An alternative strategy would be to choose different numbers of factors based on the cumulative amount of variance explained. For the sake of simplicity and as a proof of principle, in this analysis we chose to use five factors as they explained a substantial amount of the variance in expression (17.5%) without too large a multiple testing burden. The sixth factor on average would have explained 2.2% more of the variance.

### 2.3 Pathway factor and phenotype association

Association tests were performed using the linear mixed models defined in Box 1: i) between each pathway factor and chronological age, and ii) between single genes and chronological age. These models have been implemented by the lme4 package [Bates et al., 2014] in R [R Core Team, 2013]. For each phenotype a likelihood ratio test of the full model, which includes the age term, and the null model (without modelling age) was used to assess evidence for an age effect. P values produced by this analysis were assessed for significance allowing for multiple testing using a Bonferroni adjusted threshold. Permuted datasets were created which maintained the twin structure by permuting singletons, dizygotic and monozygotic twins separately and ensuring that twin pairs were kept together.

Significant associations between phenotype factors and age were further investigated to trace the particular genes within the pathway driving the signal. We report genes with a Bonferroni significant P value which accounts for the number of genes within the pathway that was tested.

### 2.4 Heritability analysis

To compute heritability, the proportion of environmental variance explained by age, and the proportion of variance explained by unique environment, we fitted the full model from Box 1. Then the genetic component to variation was estimated as twice the additional correlation of MZ twins relative to DZ twins. The environmental component to the phenotype was the sum of the contribution from the fixed age effect, the random noise term, and the shared environmental component, again estimated from the difference between MZ and DZ. Estimates of these proportions are constrained to lie between 0 and 1 inclusive.

### 2.5 Single-gene based pathway enrichment analysis

We compared the significant pathways found by our factor analysis methods to those found by looking for enrichment of single gene associations with age. Firstly we tested each gene for association with age using the methods described in Box 1 and produced a list of Bonferroni significant genes P< 0.05 (this list contained 682 differentially expressed genes). For each pathway, we applied a Fisher’s exact test to infer whether the proportion of significantly associated genes within the pathway was greater than would be expected by chance. We also investigated whether using an FDR cut-off for significant age associations would produce more significant pathways or power would be diluted by including too many false positives. When re-running the analysis using a less stringent threshold (3,487 genes were associated with age with FDR< 0.05) we found fewer significant pathways, and results correlated less well with the results of the factor based analysis (Spearman correlation of 0.36 (P=5.1×10^−7^) compared to 0.49 for Bonferroni, P=2.1 × 10^−12^). A complete list of all significant single gene age associations (FDR< 0.05, 3,487 genes), with estimate of effect size and direction, can be found in Supplementary File 1.

## 3 Results

The first stage of the analysis was to remove the effect of both known and unknown nuisance variables from the gene expression data. Using PEER software, we estimated five global factors which explained 35.7 % of the variation in the complete gene expression data. As the aim of this analysis was to find pathway specific responses to ageing, we treated these global factors as nuisance covariates and regressed these out of the data, together with batch and RNA quality which are known experimental confounders. Data were then divided into subsets of genes within 186 KEGG pathways that contained more than 10 genes with probes in our dataset. For each pathway, five factors were estimated using PEER as described above, which explained on average 17.5% of the residual variation of all genes within this pathway after removing the global factors. For the 186 KEGG pathways, this produced 930 phenotypes which were tested for association with age (see Methods for details). In total, 69 significant associations (P < 5.38×10^−5^, the Bonferroni adjusted threshold) from 57 distinct pathways were identified. The most significant 20 pathways are listed in Table 1, and a list of all 57 significant pathways can be found in the Supplementary materials (Supplementary Table S1).

**Table 1.** List of 20 pathways most significantly associated with age, together with the total number of genes in the pathway, the number of genes within pathways significantly associated with age (P < 0.05, corrected using Bonferroni for the total number of genes in the pathway), and the heritability of the pathway factor.

| KEGG.ID | Pathway | $P$ value of<br>pathway<br>factor | Number of<br>genes in<br>pathway | Number of<br>age<br>associated<br>genes | Heritability |
| --- | --- | --- | --- | --- | --- |
| 00900 | Terpenoid Backbone<br>Biosynthesis | $6.23 \times 10^{-13}$ | 13 | 6 | 0.00 |
| 00980 | Metabolism of Xenobiotics<br>by Cytochrome P450 | $6.47 \times 10^{-13}$ | 54 | 6 | 0.09 |
| 01040 | Biosynthesis of Unsaturated<br>Fatty Acids | $1.11 \times 10^{-12}$ | 17 | 6 | 0.25 |
| 00100 | Steroid Biosynthesis | $1.33 \times 10^{-12}$ | 14 | 12 | 0.41 |
| 00650 | Butanoate Metabolism | $1.51 \times 10^{-12}$ | 27 | 8 | 0.39 |
| 04146 | Peroxisome | $1.56 \times 10^{-12}$ | 64 | 17 | 0.45 |
| 00830 | Retinol Metabolism | $1.93 \times 10^{-12}$ | 48 | 6 | 0.45 |
| 00010 | Glycolysis Gluconeogenesis | $3.59 \times 10^{-12}$ | 49 | 12 | 0.42 |
| 00051 | Fructose and Mannose<br>Metabolism | $3.99 \times 10^{-12}$ | 32 | 8 | 0.32 |
| 00290 | Valine Leucine and<br>Isoleucine Biosynthesis | $1.15 \times 10^{-11}$ | 11 | 3 | 0.00 |
| 00561 | Glycerolipid Metabolism | $2.63 \times 10^{-11}$ | 38 | 6 | 0.34 |
| 00620 | Pyruvate Metabolism | $4.20 \times 10^{-11}$ | 35 | 11 | 0.37 |
| 00770 | Pantothenate and COA<br>Biosynthesis | $4.76 \times 10^{-11}$ | 16 | 4 | 0.48 |
| 00280 | Valine Leucine and<br>Isoleucine Degradation | $5.79 \times 10^{-11}$ | 35 | 10 | 0.51 |
| 00020 | Citrate Cycle TCA Cycle | $1.12 \times 10^{-10}$ | 23 | 8 | 0.43 |
| 04916 | Melanogenesis | $3.34 \times 10^{-10}$ | 93 | 10 | 0.00 |
| 04910 | Insulin Signalling Pathway | $3.70 \times 10^{-10}$ | 122 | 13 | 0.45 |
| 00565 | Ether Lipid Metabolism | $5.89 \times 10^{-10}$ | 27 | 3 | 0.00 |
| 00350 | Tyrosine Metabolism | $9.44 \times 10^{-10}$ | 32 | 4 | 0.34 |
| 00640 | Propanoate Metabolism | $1.03 \times 10^{-9}$ | 26 | 6 | 0.59 |

We also explored an alternative method for finding pathway related to ageing, looking for enrichment in the number of significantly associated genes falling into a particular pathway, analogous to the method used in the DAVID methodology [Huang et al., 2009]. This discovered a total of 7 significant pathways (Supplementary Table S2). Thus, applying factor analysis methods to discover significantly associated pathways uncovered eight times as many hits. All pathways discovered by single gene enrichment methods were also discovered using factor analysis. There is strong concordance between P values discovered by the two methods (Spearman correlation = 0.49, P= 2.1 × 10^−12^). Figure 1 shows a Q-Q plot of p values for both methods against the theoretical p values under the complete null hypothesis. We see enrichment of significant P values for both methods, but this is not present when analysing the permuted data with factor analysis methods (green dots). This suggests that age plays a widespread role in the expression of these pathways.

**Figure 1.**
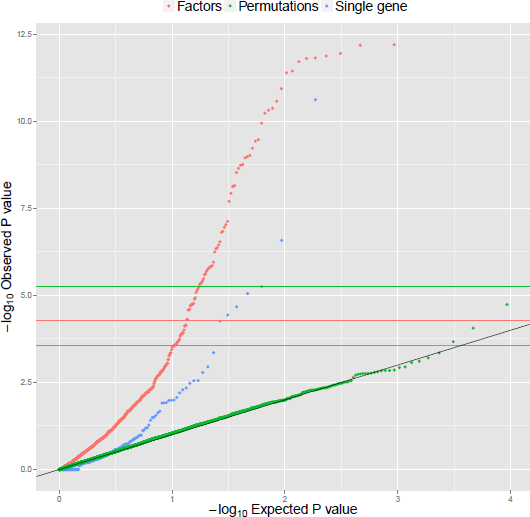
Q-Q plot of observed p values against theoretical p values for factor analysis (red dots) and single-gene based methods (in blue). Permutations (in green) shows the results of a combined analysis of 10 permuted datasets. Horizontal lines show Bonferroni significance thresholds accounting for different numbers of tests (186 tests for single gene measures in blue, 930 for factor analysis in red, and 9300 for the combined 10 permutation analyses in green).

To investigate which genes drove the significant pathway associations, we examined how many genes within a significant pathway showed significant age associations (Table 1 and Supplementary Table S1). On average 16% of genes within the pathways have P < 0.05 after adjusting for the number of genes in the pathway, with a minimum of 1 gene and maximum of 24. The proportion is similar between pathways of different sizes, in contrary to the traditional pathway enrichment analysis, where there is bias towards large pathways.

Different KEGG pathways can contain overlapping sets of genes, as they can describe related biological function. Because of this, our significant associations with age for different pathways could be related due to a common underlying effect on a given set of genes. To explore whether the observed age-associations are unique to their pathway, or common to multiple pathways, we calculated the Spearman correlation between those phenotypes. There are 24 pathway phenotypes with a correlation greater than 0.8 with at least one other phenotype (Supplementary Table S3). These phenotypes frequently relate to metabolism, and form a highly connected set (Supplementary Figure S1). We infer from this that there could be a common effect of age acting on these phenotype factors. However, these form only a minority of the phenotype factors with significant signal.

We next explored how different sources of variation in the different phenotypes analysed here affect our ability to discover age associations. We calculated the heritabilities, the proportion of environmental variance explained by age, and the proportion of variance explained by the unique environment (Box 1) for i) KEGG pathways, ii) global factors (which we have treated as nuisance covariates) and iii) for individual genes (Figure 2, global factor histograms are not shown as there are too few phenotypes). The relative differences in sources of variation between global and pathway factors, and individual genes are shown in Figure 3. We see that as we move away from local phenotypes (individual genes) to pathway phenotypes and then to global phenotypes, the proportion of variation explained by unique environment decreases. This is because that there is a stochastic component to each single gene’s expression: by taking a weighted average of a number of genes, we average away this component. If all else were to remain constant, this reduction in stochastic noise would simultaneously increase heritability (as the total variance decreases), and boost the ability to discover associations with biological meaning, such as age. We see in the first panel of Figure 3 that the relative contribution of unique environment to pathway phenotypes is smaller than the contribution to genes. This also partly explains the results shown in the second and third panels: a greater proportion of variance is explained by age and genetic factors (heritability) for pathway factors than individual gene measurements.

**Figure 2.**
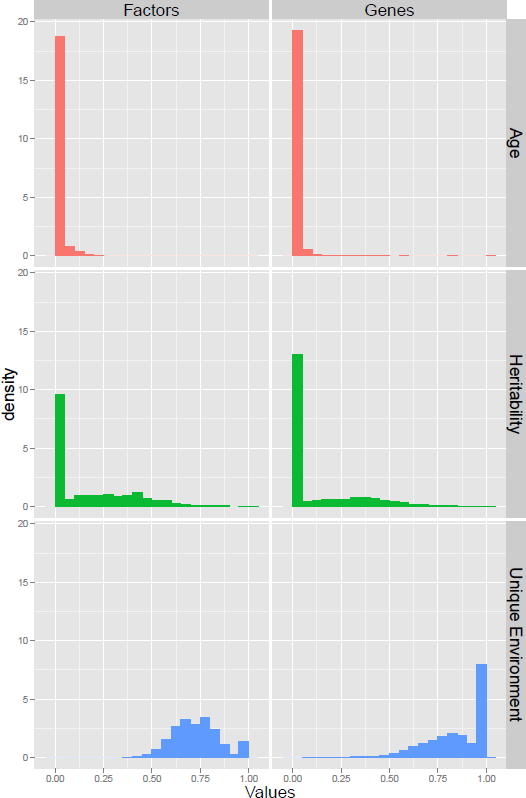
Histograms showing the proportion of environmental variation explained by age, heritability, and the proportion of variance explained by the unique environment for pathway factors and the individual gene measurements.

**Figure 3.**
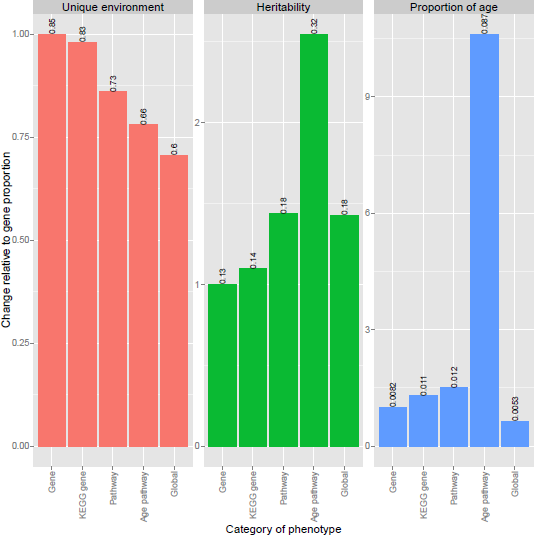
The relative importance of sources of variation to global, pathway and gene phenotypes. Measures of variation shown are the proportion of variance explained by unique environment, proportion of variance explained by genetics (heritability) and the proportion of environmental variation explained by age. To show more clearly the differences in relative importance of these measures to different classes of phenotypes, all proportions are scaled such that contribution to gene phenotypes equals one. Numbers above the bars give the absolute, unscaled proportions.

When considering global factors, as expected the unique environment is greatly reduced. However, there is not a strong influence of ageing and heritability in this case is still moderate. This is likely because age and genetics do not act in a consistent way across large sets of genes. Leek and Storey [2007] argued that global factors can capture experimental noise and batch effects. This is consistent with our findings. Heritabilities and proportion of variance explained by age for each pathway are reported in Supplementary Table S4.

We further looked for novel genetic associations with these pathway phenotypes, not seen as single gene expression associations. However, this was unsuccessful despite the increased heritability in pathway factors. This is likely due to the genetic architecture of gene regulation. Genes are regulated both in cis, where a nearby variant effects the expression of a single gene, and in trans, where a long range regulatory effect can hit multiple genes [Grundberg et al., 2012]. The genetics of pathway phenotypes is a combination of cis effects on individual genes and trans effects, potentially affecting multiple genes in the pathway. However, trans variants typically have much smaller effect size: the increase in the reliability of pathway phenotypes is insufficient to compensate for the lower power to discover trans effects. Thus, the only associations discovered were when single genes loaded heavily enough on a pathway to indirectly reflect a cis association that could be detected in a single gene test.

## 4. Discussion

We have seen that both the heritability and the proportion of environmental variance explained by age is greater for pathway phenotypes than for individual genes. Consistent with this, we found a greater proportion of associations for the pathway phenotypes than using single gene tests using this same dataset [Glass et al., 2013] (23% compared to 7% of phenotypes are significantly associated with age when using the same 0.05 FDR threshold adopted in that paper). This can be explained by our findings on the influence of unique environment on pathway phenotypes relative to single genes.

Stochasticity in gene expression, which contributes to the unique environment component that we measure, has been seen to increase with age. For example, animal model studies [Bahar et al., 2006, Herndon et al., 2002] have reported increased cell-to-cell variation in gene expression with age and tissue specific decline of functions associated to stochastic events. Others have found genes associated with longevity to be strongly regulated in older animals with low levels of stochasticity and higher levels of heritability [McCarroll et al., 2004, Viñuela et al., 2012]. The aim of our analysis was to find mean effects, rather than variance effects (though both effects are often seen together). By reducing the unique environment variable component using pathway factor analysis methods, we arguably focus much more on systematic longevity changes with age rather than the environmental stochasticity. However, it is difficult to make inference about causality with gene expression: we cannot know whether we are observing changes in expression which are driving the ageing process, or markers for it. Previous studies have suggested that the latter may be the case, as often changes in gene expression occur in response to ageing [de Magalhaes et al., 2009].

Of the 57 significant pathways, we frequently see four types of pathway, all of which have been previously linked with ageing: i) insulin signalling; ii) sugar and fatty acid metabolism; iii) xenobiotic metabolism; and iv) cancer related pathways.

We find the insulin signalling pathway (hsa04910) to be highly associated with age in our data (P = 3.7×10^−10^). Much evidence has accumulated for the influence of the insulin signalling pathway on longevity, originating in C. elegans, where lowered insulin/IGF-1 signalling (IIS) can lead to a significant increase in life span [Friedman and Johnson, 1988]. This effect has also been seen in the fruit fly D. melanogaster [Clancy et al., 2001] and in mice [Holzenberger et al., 2003]. Outside of model organisms, it has been observed that variants in FOXO transcription factors related to this pathway can affect longevity in humans [Willcox et al., 2008].

In addition to those related to insulin, our list of age-associated pathways includes many that are involved in metabolism or glycolosis. Examples of these include biosynthesis of unsaturated fatty acids (hsa00980), butanoate metabolism (hsa00650), glycolysis gluconeogenesis (hsa00010), fructose and mannose metabolism (hsa00051) and valine leucine and isoleucine biosynthesis (hsa00290). It has previously been suggested that metabolism related pathways play roles in ageing and ageing related diseases [Barzilai et al., 2012]. In particular, Houtkooper et al. [2011] showed that glucose and compounds involved in the metabolism of glucose were biomarkers of ageing in liver and muscle tissue in mice.

Other ageing related pathways include those involved in the metabolism of xenobiotics that allow cells to deactivate and excrete unexpected compounds. One example is glutathione metabolism (hsa00480, P = 1.45×10^−7^); glutathione is a well known anti-oxidant which protects against cell damage by reactive oxygen species [Pompella et al., 2003].

Finally, previous studies have shown that cancer risk is positively associated with age after childhood [Finkel et al., 2007, de Magalhães, 2013]. For example, cellular senescence, when a cell loses the ability to divide, can form a break on cancer development, and clearing such senescent cells can delay the development of age-associated disorders [Baker et al., 2011]. There are a number of pathways in our list that have been linked to cancer, in particular skin cancer. These include melanogenesis (hsa04916, P = 3.34 × 10^−10^), the PPAR signalling pathway (hsa03320, P = 1.83 × 10^−9^), the hedgehog signalling pathway (hsa04340, P = 1.12 × 10^−7^) and glioma (hsa05214, P = 4.26 × 10^−7^)

In addition to age, other phenotypes have been linked to expression patterns of multiple genes. For example, BMI has been linked to expression patterns in adipose tissue of multiple genes within a group which share a common trans master regulator, and such phenotypes could mediate between expression and diseases such as type 2 diabetes [Small et al., 2011]. Principal components and factor analysis have also been suggested as a way to build classifiers for binary traits [Hastie et al., 2000], perhaps to predict prognosis of disease from gene expression data. The ability of pathway phenotypes to provide reliable measures of expression with direct biological interpretation means they could also be applied in these situations, to understand the relationship between expression and such phenotypes.

Our analysis shows that factor analysis applied to gene expression data effectively reduces stochastic noise in summaries of gene expression patterns, giving more power to discover associations. These phenotypes are substantially more heritable than individual genes. Using them we can improve our ability to identify biological processes underpinning ageing. This is consistent with the idea that removing latent factors that exert broad effects on gene expressions increases power in associations. We show that the same idea can be used to create pathway factors that are robust and interpretable. Finally, our analysis reveals pathways that have been seen to be important in longevity from a number of previous studies, as well as novel pathways that can be further investigated.

## 5. Acknowledgement

We thank Silvia Chiappa and David A. Knowles for helpful discussions regarding the models. We thank the twins for their voluntary contribution to this project. Andrew Brown and Ana Viñuela are supported by the EU FP7 grant EuroBATS (No. 259749). In addition, Andrew Brown is supported by a grant from the South-Eastern Norway Health Authority, (No. 2011060). Richard Durbin is supported by Wellcome Trust grant WT098051. The data are derived from samples from the TwinsUK cohort, which is funded by the Wellcome Trust and the European Community’s Seventh Framework Programme (FP7/2007-2013). TwinsUK also receives support from the National Institute for Health Research (NIHR) Clinical Research Facility at Guy’s and St Thomas’ NHS Foundation Trust and NIHR Biomedical Research Centre based at Guy’s and St Thomas’ NHS Foundation Trust and King’s College London. Tim Spector is an NIHR Senior Investigator and is holder of an ERC Advanced Principal Investigator award.

---

**Table S1.** List of all pathways significantly associated with age, together with the number of genes significantly associated with age (p < 0.05, corrected using Bonferroni for the total number of genes in the pathway) and the total number of genes in the pathway.

| KEGG ID | Pathway | $P$ value | Number of<br>age<br>associated<br>genes | Total number<br>of genes |
| --- | --- | --- | --- | --- |
| 00900 | Terpenoid Backbone Biosynthesis | $6.23 \times 10^{-13}$ | 6 | 13 |
| 00980 | Metabolism of Xenobiotics By<br>Cytochrome P450 | $6.47 \times 10^{-13}$ | 6 | 54 |
| 01040 | Biosynthesis of Unsaturated Fatty<br>Acids | $1.11 \times 10^{-12}$ | 6 | 17 |
| 00100 | Steroid Biosynthesis | $1.33 \times 10^{-12}$ | 12 | 14 |
| 00650 | Butanoate Metabolism | $1.51 \times 10^{-12}$ | 8 | 27 |
| 04146 | Peroxisome | $1.56 \times 10^{-12}$ | 17 | 64 |
| 00830 | Retinol Metabolism | $1.93 \times 10^{-12}$ | 6 | 48 |
| 00010 | Glycolysis Gluconeogenesis | $3.59 \times 10^{-12}$ | 12 | 49 |
| 00051 | Fructose and Mannose Metabolism | $3.99 \times 10^{-12}$ | 8 | 32 |
| 00290 | Valine Leucine and Isoleucine<br>Biosynthesis | $1.15 \times 10^{-11}$ | 3 | 11 |
| 00561 | Glycerolipid Metabolism | $2.63 \times 10^{-11}$ | 6 | 38 |
| 00620 | Pyruvate Metabolism | $4.2 \times 10^{-11}$ | 11 | 35 |
| 00770 | Pantothenate and COA<br>Biosynthesis | $4.76 \times 10^{-11}$ | 4 | 16 |
| 00280 | Valine Leucine and Isoleucine<br>Degradation | $5.79 \times 10^{-11}$ | 10 | 35 |
| 00020 | Citrate Cycle TCA Cycle | $1.12 \times 10^{-10}$ | 8 | 23 |
| 04916 | Melanogenesis | $3.34 \times 10^{-10}$ | 10 | 93 |

| KEGG ID | Pathway | *P* value | Age-associated genes | Total number of genes |
| --- | --- | --- | --- | --- |
| 04910 | Insulin Signalling Pathway | $3.7 \times 10^{-10}$ | 13 | 122 |
| 00565 | Ether Lipid Metabolism | $5.89 \times 10^{-10}$ | 3 | 27 |
| 00350 | Tyrosine Metabolism | $9.44 \times 10^{-10}$ | 4 | 32 |
| 00640 | Propanoate Metabolism | $1.03 \times 10^{-9}$ | 6 | 26 |
| 04530 | Tight Junction | $1.12 \times 10^{-9}$ | 11 | 106 |
| 00030 | Pentose Phosphate Pathway | $1.74 \times 10^{-9}$ | 8 | 21 |
| 03320 | PPAR Signalling Pathway | $1.83 \times 10^{-9}$ | 10 | 56 |
| 00630 | Glyoxylate and Dicarboxylate Metabolism | $2.22 \times 10^{-9}$ | 4 | 11 |
| 00982 | Drug Metabolism Cytochrome P450 | $2.93 \times 10^{-9}$ | 6 | 55 |
| 00260 | Glycine Serine and Threonine Metabolism | $7.02 \times 10^{-9}$ | 4 | 30 |
| 00140 | Steroid Hormone Biosynthesis | $7.49 \times 10^{-9}$ | 7 | 44 |
| 00380 | Tryptophan Metabolism | $1.17 \times 10^{-8}$ | 6 | 32 |
| 04930 | Type II Diabetes Mellitus | $1.98 \times 10^{-8}$ | 5 | 44 |
| 05412 | Arrhythmogenic Right Ventricular Cardiomyopathy Arvc | $7.44 \times 10^{-8}$ | 7 | 70 |
| 00052 | Galactose Metabolism | $9.27 \times 10^{-8}$ | 3 | 24 |
| 04340 | Hedgehog Signaling Pathway | $1.12 \times 10^{-7}$ | 7 | 52 |
| 00480 | Glutathione Metabolism | $1.45 \times 10^{-7}$ | 7 | 39 |
| 00532 | Glycosaminoglycan Biosynthesis Chondroitin Sulfate | $1.53 \times 10^{-7}$ | 5 | 16 |
| 04920 | Adipocytokine Signaling Pathway | $2.87 \times 10^{-7}$ | 9 | 61 |
| 05214 | Glioma | $4.26 \times 10^{-7}$ | 6 | 59 |
| 05322 | Systemic Lupus Erythematosus | $4.56 \times 10^{-7}$ | 7 | 87 |
| 05414 | Dilated Cardiomyopathy | $5.64 \times 10^{-7}$ | 6 | 84 |

| KEGG ID | Pathway | *P* value | Age-associated genes | Total number of genes |
| --- | --- | --- | --- | --- |
| 00410 | Beta Alanine Metabolism | $1.11 \times 10^{-6}$ | 4 | 19 |
| 00330 | Arginine and Proline Metabolism | $1.39 \times 10^{-6}$ | 11 | 47 |
| 04510 | Focal Adhesion | $1.47 \times 10^{-6}$ | 18 | 173 |
| 00340 | Histidine Metabolism | $1.53 \times 10^{-6}$ | 3 | 25 |
| 04360 | Axon Guidance | $1.66 \times 10^{-6}$ | 15 | 119 |
| 04060 | ECM Receptor Interaction | $1.77 \times 10^{-6}$ | 13 | 71 |
| 04150 | MTOR Signaling Pathway | $2.02 \times 10^{-6}$ | 3 | 43 |
| 04270 | Vascular Smooth Muscle Contraction | $3.31 \times 10^{-6}$ | 14 | 103 |
| 00071 | Fatty Acid Metabolism | $3.84 \times 10^{-6}$ | 8 | 30 |
| 04142 | Lysosome | $4.43 \times 10^{-6}$ | 14 | 106 |
| 00983 | Drug Metabolism Other Enzymes | $5.71 \times 10^{-6}$ | 4 | 43 |
| 00040 | Pentose and Glucuronate Interconversions | $6.49 \times 10^{-6}$ | 1 | 21 |
| 05416 | Viral Myocarditis | $1.16 \times 10^{-5}$ | 5 | 51 |
| 00520 | Amino Sugar and Nucleotide Sugar Metabolism | $1.7 \times 10^{-5}$ | 7 | 39 |
| 05217 | Basal Cell Carcinoma | $1.8 \times 10^{-5}$ | 10 | 52 |
| 00510 | N-Glycan Biosynthesis | $1.82 \times 10^{-5}$ | 7 | 40 |
| 04260 | Cardiac Muscle Contraction | $1.83 \times 10^{-5}$ | 5 | 59 |
| 05216 | Thyroid Cancer | $1.99 \times 10^{-5}$ | 8 | 60 |
| 05120 | Epithelial Cell Signaling in Helicobacter Pylori Infection | $4.85 \times 10^{-5}$ | 11 | 59 |

**Table S2.**
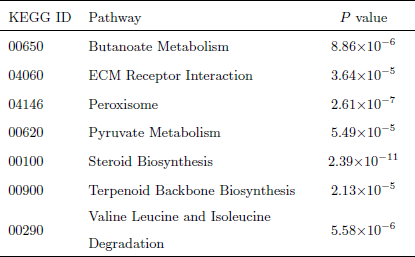
List of the seven pathways which were significantly associated with age, discovered by looking for enrichment of single gene age associations.

**Figure S1.**
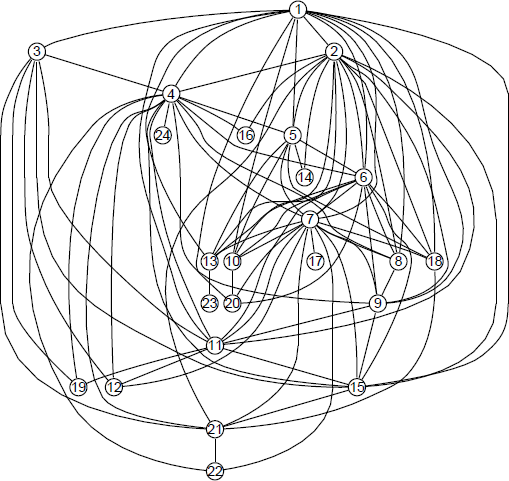
Network of connected factor phenotypes. Twenty four of the 69 age-associated factor phenotypes have a Spearman correlation of at least 0.8 with at least one other phenotype. These phenotypes show a highly connected structure, likely meaning there are common age effects driving these associations. A key for identifying which pathways correspond to the nodes can be found in Supplementary Table S3.

**Table S3.**
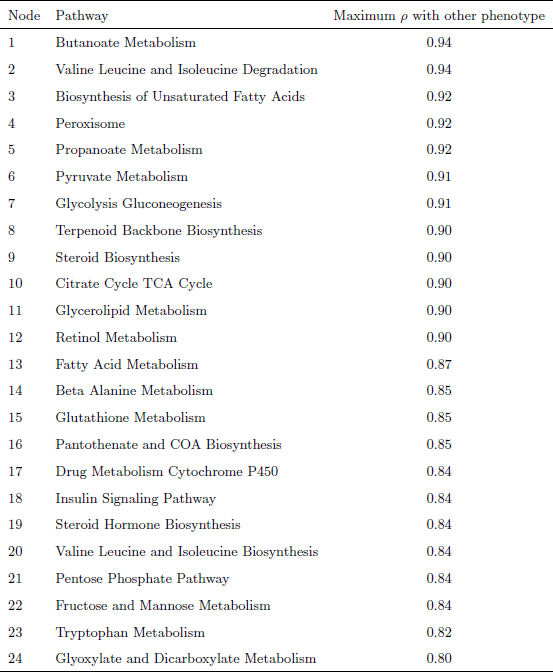
Key showing which pathways correspond to which nodes in Supplementary Figure S1, and the maximum Spearman correlation of that phenotype with any of the others representing pathways.

**Table S4.** Heritability and proportion of environmental variation explained by age for all pathways. Value reported is for the pathway phenotype most significantly associated with ageing.

| KEGG ID | Pathway | Heritability | Proportion<br>(age) |
| --- | --- | --- | --- |
| 00900 | Terpenoid Backbone Biosynthesis | $1.53 \times 10^{-11}$ | 0.0898 |
| 00980 | Metabolism of Xenobiotics By<br>Cytochrome P450 | 0.0904 | 0.0986 |
| 01040 | Biosynthesis of Unsaturated Fatty<br>Acids | 0.253 | 0.11 |
| 00100 | Steroid Biosynthesis | 0.406 | 0.143 |
| 00650 | Butanoate Metabolism | 0.39 | 0.137 |
| 04146 | Peroxisome | 0.453 | 0.152 |
| 00830 | Retinol Metabolism | 0.449 | 0.149 |
| 00010 | Glycolysis Gluconeogenesis | 0.417 | 0.14 |
| 00051 | Fructose and Mannose Metabolism | 0.316 | 0.109 |
| 00290 | Valine Leucine and Isoleucine<br>Biosynthesis | $2.61 \times 10^{-12}$ | 0.0771 |
| 00561 | Glycerolipid Metabolism | 0.337 | 0.113 |
| 00620 | Pyruvate Metabolism | 0.368 | 0.117 |
| 00770 | Pantothenate and COA<br>Biosynthesis | 0.477 | 0.136 |
| 00280 | Valine Leucine and Isoleucine<br>Degradation | 0.51 | 0.147 |
| 00020 | Citrate Cycle TCA Cycle | 0.436 | 0.126 |
| 04916 | Melanogenesis | $2.23 \times 10^{-16}$ | 0.0708 |
| 04910 | Insulin Signaling Pathway | 0.453 | 0.121 |
| 00565 | Ether Lipid Metabolism | $1.13 \times 10^{-15}$ | 0.064 |
| 00350 | Tyrosine Metabolism | 0.342 | 0.0975 |
| 00640 | Propanoate Metabolism | 0.591 | 0.157 |

| KEGG ID | Pathway | Heritability | Proportion |
| --- | --- | --- | --- |
| 04530 | Tight Junction | 0.103 | 0.0751 |
| 00030 | Pentose Phosphate Pathway | 0.291 | 0.0831 |
| 03320 | PPAR Signaling Pathway | 0.235 | 0.0777 |
| 00630 | Glyoxylate and Dicarboxylate Metabolism | 0.275 | 0.0836 |
| 00982 | Drug Metabolism Cytochrome P450 | 0.248 | 0.0811 |
| 00260 | Glycine Serine and Threonine Metabolism | 0.599 | 0.141 |
| 00140 | Steroid Hormone Biosynthesis | 0.655 | 0.167 |
| 00380 | Tryptophan Metabolism | 0 | 0.0491 |
| 04930 | Type II Diabetes Mellitus | 0.594 | 0.13 |
| 05412 | Arrhythmogenic Right Ventricular Cardiomyopathy Arvc | 0.241 | 0.0674 |
| 00052 | Galactose Metabolism | $3.4 \times 10^{-11}$ | 0.0504 |
| 04340 | Hedgehog Signaling Pathway | 0.375 | 0.08 |
| 00480 | Glutathione Metabolism | 0.415 | 0.0804 |
| 00532 | Glycosaminoglycan Biosynthesis Chondroitin Sulfate | 0.273 | 0.0682 |
| 04920 | Adipocytokine Signaling Pathway | $1.3 \times 10^{-20}$ | 0.0475 |
| 05214 | Glioma | 0.102 | 0.0466 |
| 05322 | Systemic Lupus Erythematosus | $8.17 \times 10^{-17}$ | 0.045 |
| 05414 | Dilated Cardiomyopathy | 0.532 | 0.0867 |
| 00410 | Beta Alanine Metabolism | 0.709 | 0.14 |
| 00330 | Arginine and Proline Metabolism | $1.7 \times 10^{-16}$ | 0.0402 |
| 04510 | Focal Adhesion | 0.397 | 0.0669 |
| 00340 | Histidine Metabolism | 0.519 | 0.0874 |
| 04360 | Axon Guidance | 0.606 | 0.0995 |
| 04060 | ECM Receptor Interaction | 0.792 | 0.196 |

| KEGG ID | Pathway | Heritability | Proportion |
| --- | --- | --- | --- |
| 04150 | MTOR Signaling Pathway | 0.219 | 0.0511 |
| 04270 | Vascular Smooth Muscle<br>Contraction | 0.27 | 0.0542 |
| 00071 | Fatty Acid Metabolism | 0.823 | 0.204 |
| 04142 | Lysosome | 0.566 | 0.0804 |
| 00983 | Drug Metabolism Other Enzymes | 0 | 0.0322 |
| 00040 | Pentose and Glucuronate<br>Interconversions | 0.562 | 0.0792 |
| 05416 | Viral Myocarditis | 0.569 | 0.0815 |
| 00520 | Amino Sugar and Nucleotide<br>Sugar Metabolism | 0.453 | 0.0577 |
| 05217 | Basal Cell Carcinoma | 0.593 | 0.0799 |
| 00510 | N Glycan Biosynthesis | $5.87 \times 10^{-16}$ | 0.0313 |
| 04260 | Cardiac Muscle Contraction | $8.3 \times 10^{-13}$ | 0.0312 |
| 05216 | Thyroid Cancer | $2.56 \times 10^{-9}$ | 0.0332 |
| 05120 | Epithelial Cell Signaling in<br>Helicobacter Pylori Infection | 0.652 | 0.0859 |
| 04060 | Cytokine Cytokine Receptor<br>Interaction | $3.51 \times 10^{-17}$ | 0.0276 |
| 00120 | Primary Bile Acid Biosynthesis | $1.69 \times 10^{-16}$ | 0.0265 |
| 00190 | Oxidative Phosphorylation | $1.41 \times 10^{-11}$ | 0.0268 |
| 00760 | Nicotinate and Nicotinamide<br>Metabolism | 0.401 | 0.0433 |
| 00360 | Phenylalanine Metabolism | 0.711 | 0.088 |
| 00512 | O Glycan Biosynthesis | $1.78 \times 10^{-18}$ | 0.0253 |
| 05213 | Endometrial Cancer | 0.428 | 0.0408 |
| 00250 | Alanine Aspartate and Glutamate<br>Metabolism | 0.526 | 0.0507 |
| 00564 | Glycerophospholipid Metabolism | 0 | 0.0231 |

| KEGG ID | Pathway | Heritability | Proportion |
| --- | --- | --- | --- |
| 04012 | ERBB Signaling Pathway | 0.121 | 0.0253 |
| 05211 | Renal Cell Carcinoma | $3.64 \times 10^{-11}$ | 0.0237 |
| 02010 | ABC Transporters | 0.506 | 0.0454 |
| 04710 | Circadian Rhythm Mammal | 0.0407 | 0.0292 |
| 05222 | Small Cell Lung Cancer | $1.03 \times 10^{-17}$ | 0.024 |
| 04062 | Chemokine Signaling Pathway | 0.124 | 0.0277 |
| 00590 | Arachidonic Acid Metabolism | 0.141 | 0.027 |
| 04610 | Complement and Coagulation<br>Cascades | 0.504 | 0.0453 |
| 03022 | Basal Transcription Factors | 0.537 | 0.0424 |
| 00600 | Sphingolipid Metabolism | $8.68 \times 10^{-19}$ | 0.0219 |
| 05410 | Hypertrophic Cardiomyopathy<br>Hcm | $3.3 \times 10^{-13}$ | 0.0147 |
| 04912 | GNRH Signaling Pathway | $3.11 \times 10^{-16}$ | 0.0187 |
| 04720 | Long Term Potentiation | 0 | 0.0183 |
| 03050 | Proteasome | 0.425 | 0.0314 |
| 04620 | JAK Stat Signaling Pathway | 0.503 | 0.0382 |
| 05330 | Allograft Rejection | 0 | 0.016 |
| 03450 | Non Homologous End Joining | 0.132 | 0.0199 |
| 05320 | Autoimmune Thyroid Disease | 0 | 0.0156 |
| 03060 | Protein Export | 0.235 | 0.0197 |
| 03420 | Nucleotide Excision Repair | $3.19 \times 10^{-14}$ | 0.0178 |
| 00660 | Alpha Linolenic Acid Metabolism | 0.458 | 0.0311 |
| 04144 | Endocytosis | 0.0714 | 0.0181 |
| 05010 | Alzheimers Disease | 0.0757 | 0.0172 |
| 00591 | Linoleic Acid Metabolism | $3 \times 10^{-11}$ | 0.0159 |
| 00240 | Pyrimidine Metabolism | $6.42 \times 10^{-13}$ | 0.0152 |

| KEGG ID | Pathway | Heritability | Proportion |
| --- | --- | --- | --- |
| 00270 | Cysteine and Methionine Metabolism | 0.00281 | 0.0162 |
| 03410 | Base Excision Repair | 0.377 | 0.0219 |
| 04722 | Neurotrophin Signaling Pathway | $4.88 \times 10^{-18}$ | 0.0152 |
| 04070 | Phosphatidylinositol Signaling System | 0.312 | 0.0207 |
| 04960 | Aldosterone Regulated Sodium Reabsorption | $3.36 \times 10^{-15}$ | 0.0142 |
| 05130 | Pathogenic Escherichia Coli Infection | 0.158 | 0.0158 |
| 04310 | WNT Signaling Pathway | 0.176 | 0.0174 |
| 00562 | Inositol Phosphate Metabolism | $3.24 \times 10^{-16}$ | 0.0138 |
| 05221 | Acute Myeloid Leukemia | 0.472 | 0.0268 |
| 00071 | Selenoamino Acid Metabolism | $3.71 \times 10^{-10}$ | 0.0137 |
| 04742 | Taste Transduction | 0.149 | 0.0174 |
| 00531 | Glycosaminoglycan Degradation | $2.23 \times 10^{-19}$ | 0.0135 |
| 05340 | Primary Immunodeficiency | 0 | 0.0133 |
| 04640 | Hematopoietic Cell Lineage | $2.35 \times 10^{-16}$ | 0.0132 |
| 05310 | Asthma | 0.331 | 0.0183 |
| 04620 | TGF Beta Signaling Pathway | $1.72 \times 10^{-18}$ | 0.0131 |
| 00860 | Porphyrin and Chlorophyll Metabolism | $9.84 \times 10^{-16}$ | 0.0124 |
| 04612 | Antigen Processing and Presentation | $2.03 \times 10^{-11}$ | 0.0129 |
| 05010 | Parkinsons Disease | $4.25 \times 10^{-9}$ | 0.012 |
| 00790 | Folate Biosynthesis | $1.07 \times 10^{-11}$ | 0.0119 |
| 00500 | Starch and Sucrose Metabolism | 0.429 | 0.0111 |
| 05223 | Non Small Cell Lung Cancer | 0 | 0.0115 |
| 03030 | DNA Replication | 0 | 0.0116 |

| KEGG ID | Pathway | Heritability | Proportion |
| --- | --- | --- | --- |
| 04622 | RIG I Like Receptor Signaling Pathway | 0 | 0.0117 |
| 04666 | FC Gamma R Mediated Phagocytosis | 0.747 | 0.0415 |
| 04514 | Cell Adhesion Molecules CAMS | 0.278 | 0.016 |
| 03430 | Mismatch Repair | $7.18 \times 10^{-17}$ | 0.011 |
| 03010 | Ribosome | $8.63 \times 10^{-19}$ | 0.0108 |
| 05220 | Chronic Myeloid Leukemia | 0.333 | 0.0164 |
| 00910 | Nitrogen Metabolism | 0 | 0.0106 |
| 04330 | Notch Signaling Pathway | 0.585 | 0.0251 |
| 04520 | Adherens Junction | $1.15 \times 10^{-9}$ | 0.0107 |
| 05210 | Colorectal Cancer | 0.289 | 0.0141 |
| 03018 | RNA Degradation | $1.03 \times 10^{-13}$ | 0.00998 |
| 03440 | Homologous Recombination | 0 | 0.0093 |
| 00920 | Sulfur Metabolism | 0.121 | 0.011 |
| 00310 | Lysine Degradation | 0.446 | 0.0166 |
| 04662 | B Cell Receptor Signaling Pathway | 0.494 | 0.0183 |
| 00430 | Taurine and Hypotaurine Metabolism | $8.53 \times 10^{-13}$ | 0.00891 |
| 04964 | Proximal Tubule Bicarbonate Reclamation | 0.456 | 0.0163 |
| 04614 | Renin Angiotensin System | 0.556 | 0.0183 |
| 00970 | Aminoacyl tRNA Biosynthesis | 0.107 | 0.0102 |
| 04672 | Intestinal Immune Network For IGA Production | 0 | 0.00883 |
| 04810 | Regulation of Actin Cytoskeleton | 0.215 | 0.0104 |
| 05215 | Prostate Cancer | $1.55 \times 10^{-9}$ | 0.00719 |
| 00563 | Glycosylphosphatidylinositol Gpi Anchor Biosynthesis | 0 | 0.00816 |

| KEGG ID | Pathway | Heritability | Proportion |
| --- | --- | --- | --- |
| 04660 | NOD Like Receptor Signaling Pathway | 0 | 0.00828 |
| 04540 | Gap Junction | 0.121 | 0.0096 |
| 00903 | Limonene and Pinene Degradation | $4.8 \times 10^{-12}$ | 0.00822 |
| 05200 | Pathways in Cancer | 0.275 | 0.0119 |
| 04660 | Toll Like Receptor Signaling Pathway | $8.13 \times 10^{-17}$ | 0.00782 |
| 04730 | Long Term Depression | 0.128 | 0.00885 |
| 04020 | Calcium Signaling Pathway | 0.148 | 0.00936 |
| 04320 | Dorso Ventral Axis Formation | 0.271 | 0.00857 |
| 05110 | Vibrio Cholerae Infection | 0.353 | 0.011 |
| 04115 | P53 Signaling Pathway | 1.07 | -0.0975 |
| 04962 | Vasopressin Regulated Water Reabsorption | 0.331 | 0.0107 |
| 04670 | Leukocyte Transendothelial Migration | 0.248 | 0.00871 |
| 03020 | RNA Polymerase | $2.52 \times 10^{-16}$ | 0.00609 |
| 04664 | FC Epsilon RI Signaling Pathway | 0.35 | 0.00908 |
| 04140 | Regulation of Autophagy | 0 | 0.00509 |
| 05010 | Huntingtons Disease | 0.894 | 0.0529 |
| 00670 | One Carbon Pool By Folate | $9.11 \times 10^{-13}$ | 0.00564 |
| 04660 | T Cell Receptor Signaling Pathway | 0.487 | 0.0103 |
| 00740 | Riboflavin Metabolism | 0.252 | 0.00627 |
| 00533 | Glycosaminoglycan Biosynthesis Keratan Sulfate | 0 | 0.00452 |
| 00230 | Purine Metabolism | $3.84 \times 10^{-18}$ | 0.00462 |
| 04130 | Snare Interactions in Vesicular Transport | $1.2 \times 10^{-17}$ | 0.00475 |
| 05020 | Prion Diseases | 0.272 | 0.0059 |

| KEGG ID | Pathway | Heritability | Proportion |
| --- | --- | --- | --- |
| 05219 | Bladder Cancer | 0.229 | 0.00531 |
| 03040 | Spliceosome | 0.224 | 0.00573 |
| 04010 | Mapk Signaling Pathway | 0.221 | 0.00506 |
| 00534 | Glycosaminoglycan Biosynthesis<br>Heparan Sulfate | $1.4 \times 10^{-18}$ | 0.00416 |
| 00604 | Glycosphingolipid Biosynthesis<br>Ganglio Series | 0 | 0.00372 |
| 04940 | Type I Diabetes Mellitus | 0.446 | 0.00735 |
| 04623 | Cytosolic DNA Sensing Pathway | 0.431 | 0.00706 |
| 05332 | Graft Versus Host Disease | 0.432 | 0.00691 |
| 04740 | Olfactory Transduction | 0 | 0.0035 |
| 04110 | Cell Cycle | $5.02 \times 10^{-18}$ | 0.00369 |
| 00511 | Other Glycan Degradation | $1.07 \times 10^{-24}$ | 0.00321 |
| 05140 | Leishmania Infection | 0.136 | 0.00381 |
| 04914 | Progesterone Mediated Oocyte<br>Maturation | $1.82 \times 10^{-19}$ | 0.00322 |
| 04120 | Ubiquitin Mediated Proteolysis | $2.55 \times 10^{-15}$ | 0.00315 |
| 00604 | Glycosphingolipid Biosynthesis<br>Globo Series | 0 | 0.00271 |
| 00601 | Glycosphingolipid Biosynthesis<br>Lacto and Neolacto Series | 0.213 | 0.00341 |
| 04370 | VEGF Signaling Pathway | 0.192 | 0.00362 |
| 00053 | Ascorbate and Aldarate<br>Metabolism | 0 | 0.00197 |
| 04650 | Natural Killer Cell Mediated<br>Cytotoxicity | $4.16 \times 10^{-19}$ | 0.00222 |
| 05212 | Pancreatic Cancer | $5.99 \times 10^{-48}$ | 0.00212 |
| 04114 | Oocyte Meiosis | $1.82 \times 10^{-11}$ | 0.00201 |
| 04210 | Apoptosis | 0.632 | 0.00523 |

| KEGG ID | Pathway | Heritability | Proportion |
| --- | --- | --- | --- |
| 05218 | Melanoma | 0.349 | 0.00284 |
| 04080 | Neuroactive Ligand Receptor<br>Interaction | $1.76 \times 10^{-17}$ | 0.00158 |
| 05014 | Amyotrophic Lateral Sclerosis ALS | 0 | 0.00102 |
| 04950 | Maturity Onset Diabetes of The<br>Young | $8.21 \times 10^{-12}$ | 0.000707 |

